# Low tolerance for transcriptional variation at cohesin genes is accompanied by functional links to disease-relevant pathways

**DOI:** 10.1101/2020.04.11.037358

**Authors:** William Schierding, Julia Horsfield, Justin O’Sullivan

## Abstract

Variants in DNA regulatory elements can alter the regulation of distant genes through spatial-regulatory connections. In humans, these spatial-regulatory connections are largely set during early development, when the cohesin complex plays an essential role in genome organisation and cell division. A full complement of the cohesin complex and its regulators is important for normal development, since heterozygous mutations in genes encoding these components are often sufficient to produce a disease phenotype. The implication that genes encoding the cohesin complex and cohesin regulators must be tightly controlled and resistant to variability in expression has not yet been formally tested. Here, we identify spatial-regulatory connections with potential to regulate expression of cohesin loci, including linking their expression to that of other genes. Connections that centre on the cohesin ring subunits (Mitotic: SMC1A, SMC3, STAG1, STAG2, RAD21/RAD21-AS; Meiotic: SMC1B, STAG3, REC8, RAD21L1), cohesin-ring support genes (NIPBL, MAU2, WAPL, PDS5A and PDS5B), and CTCF provide evidence of coordinated regulation that has little tolerance for perturbation. We identified transcriptional changes across a set of genes co-regulated with the cohesin loci that include biological pathways such as extracellular matrix production and proteasome-mediated protein degradation. Remarkably, many of the genes that are co-regulated with cohesin loci are themselves intolerant to loss-of-function. The results highlight the importance of robust regulation of cohesin genes, indicating novel pathways that may be important in the human cohesinopathy disorders.

## Introduction

The cohesin complex has multiple essential roles during cell division in mitosis and meiosis, genome organisation, DNA damage repair, and gene expression^1^. Mutations in genes that encode members of the cohesin complex, or its regulators, cause developmental diseases known as the ‘cohesinopathies’ when present in the germline^2^; or contribute to the development of cancer in somatic cells^3–5^.

Remarkably, cohesin mutations are almost always heterozygous, and result in depletion of the amount of functional cohesin without eliminating it altogether. Complete loss of cohesin is not tolerated in healthy individuals^2^. Thus, cohesin is haploinsufficient such that normal tissue development and homeostasis requires that the concentrations of cohesin and its regulatory factors remain tightly regulated.

The human mitotic cohesin ring contains four integral subunits: two structural maintenance proteins (SMC1A, SMC3), one stromalin HEAT-repeat domain subunit (STAG1 or STAG2), and one kleisin subunit (RAD21)^6^. Mutation of STAG2 has been linked to at least four tumour types (*e.g*. Ewing sarcoma, glioblastoma and melanoma^7^, and bladder carcinomas)^4^. Strikingly, mutations in cohesin components are especially prevalent in acute myeloid leukaemia^8–10^.

In meiotic cohesin, SMC1A is replaced by SMC1B; STAG1/2 by STAG3; and RAD21 by REC8 or RAD21L1^1^. Mutations in meiotic cohesin subunits are associated with infertility in men^11^, chromosome segregation errors and Primary Ovarian Insufficiency in women^12^.

Cohesin is loaded onto DNA by the SCC2/SCC4 complex (encoded by the NIPBL and MAU2 genes, respectively)^1^. Mutations in the cohesin loading factor NIPBL are associated with >65% cases of Cornelia de Lange Syndrome (CdLS). Remarkably, features associated with CdLS are observed with less than 30% depletion in NIPBL protein levels^13^.

The release of cohesin from DNA is achieved by WAPL, which opens up the interface connecting the SMC3 and RAD21 subunits. The PDS5A/PDS5B cohesin associated subunits affect this process by contacting cohesin to either maintain (with STAG1 and STAG2) or remove (with WAPL) the ring from DNA^1^.

Spatial organization and compaction of chromosomes in the nucleus involves non-random folding of DNA on different scales. The genome is segregated into active A compartments and inactive B compartments^14^, inside which further organisation occurs into topologically associating domains (TADs) interspersed with genomic regions with fewer interactions^14^. Cohesin participates in genome organisation by mediating ‘loop extrusion’ of DNA to form loops that anchor TAD boundaries. At TAD boundaries, cohesin colocalizes with the CCCTC binding factor (CTCF) to form chromatin loops between convergent CTCF binding sites. Fine-scale genomic interactions include chromatin loops that mediate promoter-enhancer contacts. Notably, the time- and tissue-specific formation of the fine-scale loops also requires cohesin.

The spatial organization of the genome is particularly dynamic and susceptible to disruption during development. For example, changes to TAD boundaries are associated with developmental disorders^15^. Furthermore, disruption of TAD boundaries by cohesin knockdown can lead to ectopic enhancer-promoter interactions that result in changes in gene expression^16^. Rewiring of the patterns of course and fine-scale chromatin interactions also contributes to cancer development^17,18^, including the generation of oncogenic chromosomal translocations^19,20^.

Disease-associated GWAS variants in non-coding DNA likely act through spatially organized hubs of regulatory control elements, each component of which contributes a small amount to the observed phenotype(s), as predicted by the omnigenic hypothesis^21,22^. Non-coding mutations at cohesin and cohesin-associated factors were found by genome wide association studies (GWAS-attributed variants) to track with multiple phenotypes^23^. However, the impact of genetic variants located within cohesin and its associated genes has not yet been investigated with respect to phenotype development.

We hypothesised that cohesin-associated pathologies can be affected by subtle, combinatorial changes in the regulation of cohesin genes caused by common genetic variants within control elements. Here, we link the 3D structure of the genome with eQTL data to determine if GWAS variants attributed to cohesin genes affect their transcription. We test cohesin gene-associated GWAS variants for regulatory connections beyond the cohesin genes (gene enrichment and regulatory hubs). We also identify all variants within each gene locus that had a previously determined *cis*-eQTL (GTEx Catalog) to the cohesin gene (eQTL-attributed variant list). As with the GWAS variants, we tested these variants for the presence of spatial-regulatory relationships involving genes outside of the locus. Only a few of these eQTL-attributed variants are currently implicated in disease pathways, but their regulatory relationships with cohesin suggest that they may be significant for cohesinopathy disorders.

## Results

### 140 genetic variants with regulatory potential are associated with cohesin loci

Mitotic cohesin genes (*SMC1A, SMC3, STAG1, STAG2*, and *RAD21*), meiotic cohesin genes (*SMC1B, STAG3, REC8*, and *RAD21L1*), cohesin support genes (*WAPL, NIPBL, PDS5A, PDS5B*, and *MAU2*) and *CTCF* were investigated to determine if they contain non-coding genetic variants (SNPs) that make contact in 3D with genes and therefore could directly affect gene expression (GWAS-attributed and eQTL-attributed; Table 1, Table S1).

A total of 106 GWAS-attributed genetic variants associated with disease were identified (methods) that mapped to a cohesin gene (49 GWAS; 50 SNPs) or cohesin-associated (49 GWAS; 56 SNPs) gene (Table S1). Twelve SNPs (blue, Table S1) are not listed in the current GTEx SNP dictionary (GTEx v8), while fourteen SNPs have virtually no variation in the GTEx per-tissue analysis (*i.e.* minor allele frequency [MAF] < 0.05; green, Table S1) and so were discarded. 80 SNPs passed all filters and were subsequently analysed using CoDeS3D^21^ (GWAS-attributed list; Table 1).

Within the GTEx catalogue, 187 eQTL-attributed variants associate with regulation of the cohesin gene set (Table S1). These variants associate with modified expression levels of cohesin genes in otherwise healthy individuals. Fifty-five of these variants had a MAF<0.05 (green, Table S1) and were filtered out of the eQTL-attributed set prior to CoDeS3D analysis (132 variants passed MAF filter; eQTL-attributed; Table 1). Only three variants were shared between the GWAS-attributed and eQTL-attributed variant lists, resulting in a total of 290 cohesin-associated variants (GWAS- and eQTL-attributed combined; Table 1), but only 209 variants pass all filters (GWAS- and eQTL-attributed combined; black, Table S1).

CoDeS3D integrates data on the 3-dimensional organisation of the genome (captured by Hi-C) and transcriptome (eQTL) associations across multiple tissue types (Table S2 and S3). We used the CoDeS3D algorithm to assign the 209 variants to hubs of regulatory/functional impacts by examining their potential to regulate other genes. Of the 209 variants, four had zero significant eQTLs, leaving 205 variants with significant eQTLs (128 eQTL-attributed, 80 GWAS-attributed, and 3 overlaps; Table S2 and S3). However, many of the 205 variants were not attributed the GWAS- or eQTL-attributed cohesin gene in the locus. Of the 205 variants with eQTLs, 106/187 (56.7%) eQTL-attributed variants and 37/106 (34.9%) GWAS-attributed variants had a physical (Hi-C detected) connection and significant eQTL with their attributed cohesin gene (140 total, 3 overlaps; Table 1).

Strikingly, most of the 106 variants attributed by GWAS studies to cohesin genes were <u>not</u> confirmed by spatial connection, with only 34% being *cis*-eQTLs for the attributed cohesin gene (37 of 106, 34.9%). After the CoDeS3D analysis, six of the cohesin genes have no GWAS-attributed SNPs with a regulatory connection (*STAG2, NIPBL, CTCF, SMC3, REC8*, and *RAD21*-*AS1*). Therefore, the majority of GWAS variants tested in proximity to cohesin loci have regulatory effects elsewhere in the genome. Remarkably, despite five SNPs being attributed to NIPBL by GWAS, none of these were attributed to regulation of NIPBL by our spatial eQTL analysis. Of the cohesin or cohesin-associated genes with any GWAS-attributed variants with *cis*-eQTLS (*RAD21, RAD21L1, SMC1A, SMC1B, STAG1, STAG3, MAU2, PDS5A, PDS5B*, and *WAPL*), only the *STAG1, MAU2*, and *PDS5B* loci contain more than two variants with confirmed *cis*-eQTLs. Therefore, even those loci with confirmed variant-gene GWAS attributions have very few variants with evidence of *cis*-eQTLs.

To further characterize the potential for the 209 cohesin-associated variants to alter gene regulation, we analysed histone marks, DNAse accessibility, and protein binding motifs (Haploreg v 4.1) at each location (Table S4)^24^. Most variants reside within accessible chromatin (DNAse: 61.3%) and almost all (93.1%) have at least one of three histone marks that are consistent with putative regulatory activity (promoter, 82.8%; enhancer, 45.1%; protein binding sites, 86.8%). Intriguingly, Haploreg motif prediction identified 16 of the 209 variants (7 different loci: *MAU2, PDS5B, REC8, SMC1B, STAG3, RAD21L1, STAG1*) as residing within protein binding domains associated with cohesin-related DNA interactions (*i.e.* RAD21, SMC3, and CTCF). Therefore, most of the variants like in regions associated with chromatin marks that highlight putative regulatory capabilities.

CoDeS3D predicted 140 out of 209 variants to have significant regulatory activity. We compared this to alternative functional variant prediction methods. The DeepSEA algorithm, which predicts the chromatin effects of sequence alterations by analysing the epigenetic state of a sequence, identified 28 of 209 the variants as having functional significance (<0.05, Table S5). PredictSNP2, which estimates noncoding variant classification (deleterious or neutral) from five separate prediction tools (CADD, DANN, FAT, FUN, GWAVA tools), identified 9 of 209 variants as deleterious (Table S6). Therefore, only 31/209 variants have putative functional significance predicted by these tools (28 DeepSEA, 9 PredictSNP, 6 overlaps). Therefore, these 31 variants have support from multiple methods, suggesting a potential for higher regulatory effects, and that the contrast with the Haploreg chromatin marks and GTEx measured eQTLs possibly indicates a heavy weighting against false positives in these prediction methods.

In summary, GWAS-attributed SNPs are enriched for chromatin marks (regulatory potential). However, fewer than half the SNPs in proximity to the cohesin and cohesin-associated genes physically connected with the cohesin genes they are predicted to regulate, suggesting that cohesin genes are not the direct targets of these regulatory variants.

### Pathway enrichment implicates coordinated regulation of cohesin with essential cell cycle genes

CoDeS3D identified 140 variants as being physically connected to, and associated with the expression levels of 310 genes (243 genes from eQTL-attributed variants, 141 from GWAS-attributed variants, and 74 overlap) across 6,795 significant tissue-specific regulatory connections (FDR p<0.05). Physical connections comprised 6,570 fine-scale connections (*cis*, <1Mb from the variant), 42 coarse-scale connections (*trans*-intrachromosomal, >1Mb), and 183 connections on a different chromosome (*trans*-interchromosomal) (Fig 1; Tables S2, S3). Of note, there is one cohesin-to-cohesin regulatory connection: *rs111444407*, a GWAS-associated variant (bipolar disorder) located within the *MAU2* locus has a significant *trans*-interchromosomal eQTL with *REC8*. The gene overlaps between the GWAS- and eQTL-attributed analyses are also intriguing. For example, variants in the GWAS-attributed and eQTL-attributed lists, each from a different chromosomal location (REC8 and PDS5B), modify TCF7L1 expression. As TCF7L1 is part of the Wnt pathway and is highly expressed in ovaries in the GTEx catalogue, it is notable that this gene is regulated from variants in the REC8 locus (meiosis-specific cohesin). Their co-regulatory relationships exemplify the systems of genome-regulatory hubs, with a total 74 genes overlapping the GWAS- and eQTL-attributed analyses.

**Figure 1.**
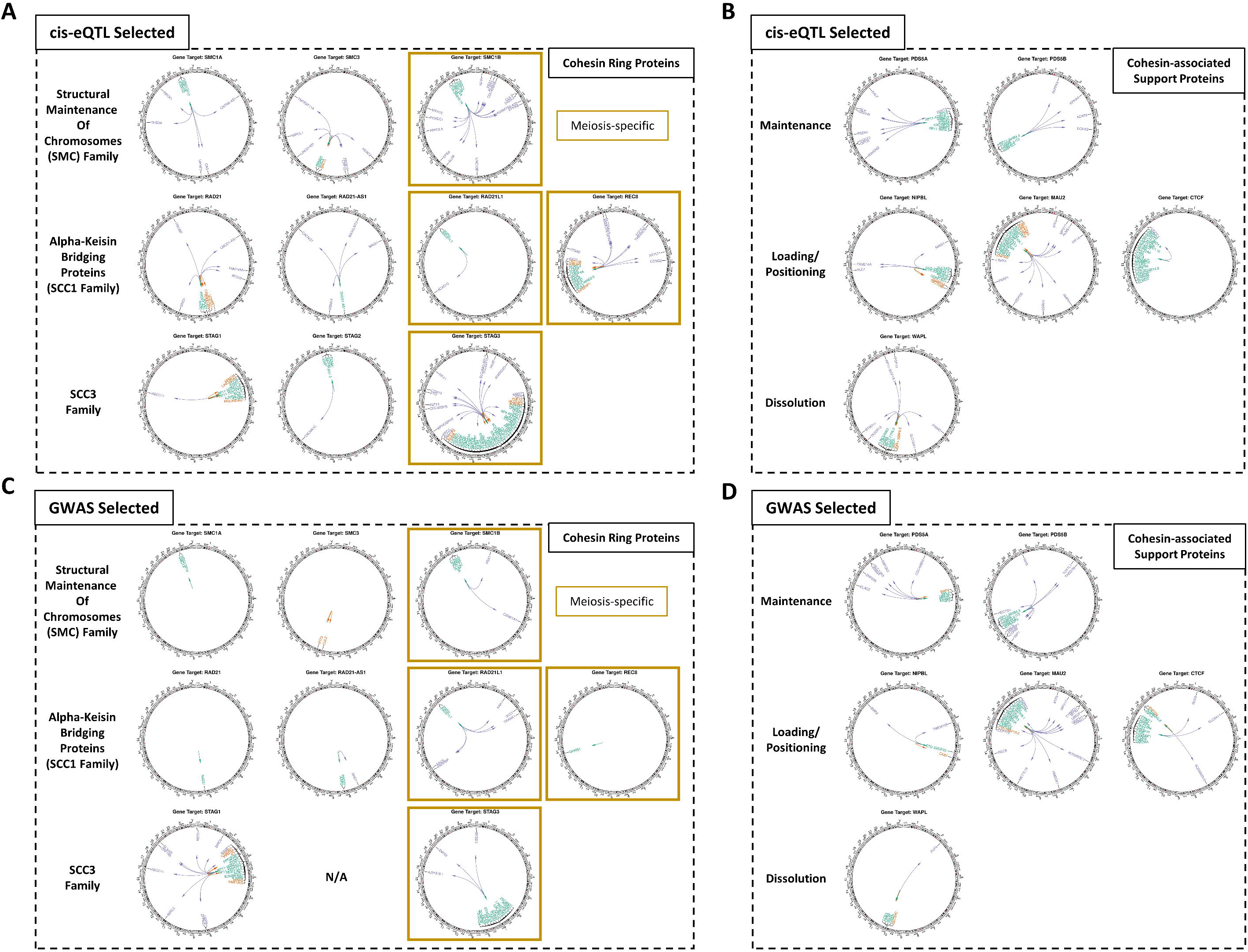
Circos plots highlighting the eQTL- and GWAS-attributed variant locations and their long-distance gene connections. Each cohesin gene has a unique set of regulatory connections across each attributed variant set (For all connections, see Supplementary Tables 2 and 3).

The 31 significant variants highlighted by the DeepSEA and PredictSNP2 analyses functionally connected to 107 (34.5%) of the genes identified by CoDeS3D. Thus, while DeepSEA and PredictSNP2 assigned functionality to just 31/205 SNPs (15.1%), these variants represent 34.5% of the CoDeS3D-predicted modulatory connections. Therefore, DeepSEA and PredictSNP2 successfully selected for variants with highly enriched regulatory functions.

We used g:Profiler to assess the functional enrichment of the GWAS- and eQTL-attributed genes (Table S7). The 310 genes are enriched for pathways that support cohesin function, as we currently understand it, within the nucleus. For example, functional enrichment in sister chromatid gene ontology categories includes five non-cohesin genes from our analysis: *CTNNB1, PPP2R1A, CHMP4A, CUL3*, and *DIS3L2*. Moreover, cohesin’s meiosis-specific role (*SMC1B, STAG3, REC8*) is enriched by two *trans* connections revealing regulation of meiosis-related genes (*ITPR2, PPP2R1A*; KEGG pathway hsa04114). Collectively, these results suggest that expression of cohesin genes is coordinated with other genes that are involved in cell cycle control.

### Meiosis-specific cohesin genes are functionally connected to KIF6 and a germ cell pathway

There are 108 genes functionally connected to variants within the meiosis-specific cohesin loci *SMC1B, STAG3, REC8*, and *RAD21L1* (Table S8a). We identified 14 significant enrichment terms using g:Profiler (Table S8b), including the gene ontology “male germ cell nucleus” pathway. The gene ontology “male germ cell nucleus” pathway contains KIF6, STAG3, and REC8 (*trans*-interchromosomal connection from STAG3 locus to KIF6). Within GTEx, *KIF6* is highly expressed in brain and testis. The *KIF6* gene region was previously identified as significantly associated with a GWAS of hypospadias, a birth defect presenting with a urethral opening located on the ventral side of the penis instead of at the tip of the glans. Of particular relevance to the cohesinopathies, in which affected individuals present with cognitive defects, a mutation in human KIF6 caused neurodevelopmental defects and intellectual disability^25^. Variation in KIF6 expression has been associated with epilepsy^26^, another notable cohesinopathy phenotype^27^. It has also been proposed that KIF6 is a conserved regulator of neurological development^25^. Previous findings of an association between KIF6 and heart disease are not supported by GTEx (*KIF6* is lowly expressed in GTEx heart tissue) and KIF6 knockout mice have no heart phenotype^28^, suggesting that the heart-associated KIF6 variants might somehow affect the expression or function of other genes.

We also identified an enrichment for E2F7 transcription factor binding sites within our genes (Table S8b). The E2F7 transcription factor modulates embryonic development and cell cycle^29,30^, with a role in cancer development^31^. Of note, the E2F7 gene is loss-of-function intolerant (pLI=.993), consistent with its crucial role in the cell cycle and development.

Collectively, our results suggest that the 301 genes are enriched for cell cycle-regulated genes, including E2F targets, and that these genes might be indirect targets of cancer drug treatments modifying E2F7 transcription factor activity.

### The cohesin gene regulatory network is intolerant to loss-of-function mutations

A subset of human genes, whose activity is crucial to survival, are intolerant to loss-of-function (LoF-intolerant) mutations^32^. The gnomAD catalog lists 19,704 genes: 3,063 LoF-intolerant, 16,134 LoF-tolerant, and 507 undetermined (15.5% LoF-intolerance, defined as pLI ≥ 0.9)^32^. All cohesin and cohesin-associated genes (except the meiosis-only *RAD21L1, SMC1B, STAG3*, and *REC8*) are LoF-intolerant (Table 2).

We hypothesized that genes functionally connecting to variants in cohesin genes would also be enriched for loss-of-function intolerance. Consistent with our hypothesis, 79 of the 310 genes (25.4%) we identified are LoF-intolerant (185 are LoF tolerant and 46 have undetermined pLI; Table S9). Stratification of the eQTL genes based on the distance between the variant and gene (*cis* versus *trans*-acting eQTL gene lists) identified a marked increase in LoF-intolerance for genes regulated by *trans*-acting eQTLs (Fig 2; Fig S1).This was especially pronounced for regulatory interactions that occurred between chromosomes (Fig 2; Fig S1). There was a significant correlation between distance and pLI on the same chromosome (r=-0.03, p<0.05).

**Figure 2.**
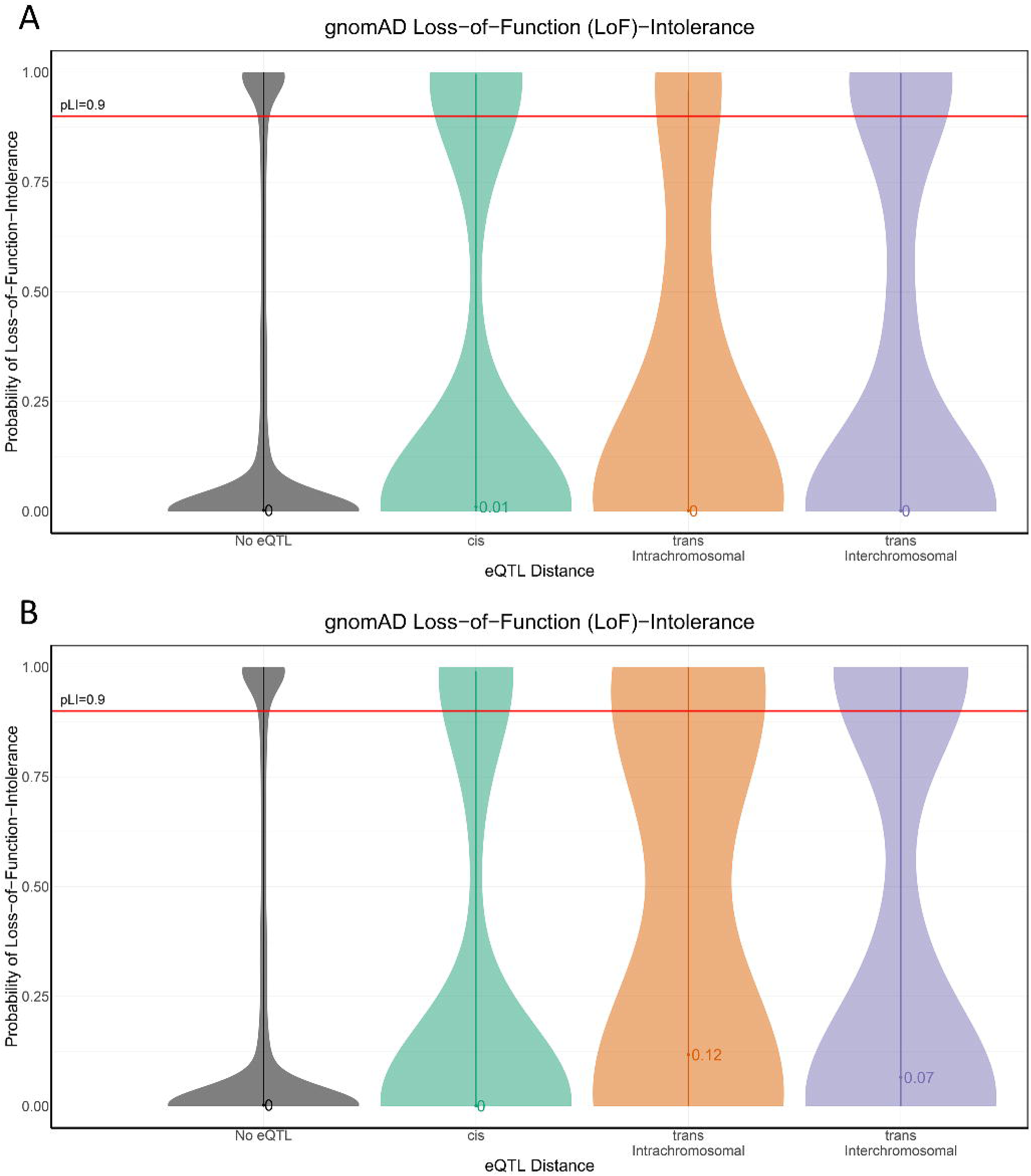
Loss-of-function intolerance shows enrichment for variant-gene eQTL distance. Violin plots (and median pLI value) of pLI across the (a) eQTL-attributed and (b) GWAS-attributed gene sets shows the changing distributions of pLI, which is increased in genes further away (*trans*) versus those genes in background (No eQTL) or nearby (*cis*).

As the cohesin genes are intolerant to even small perturbations in gene expression, we hypothesized that the LoF-intolerant genes that were regulated by elements within the cohesin genes would similarly only be tolerant to small allelic fold changes in gene expression. Therefore, we tested for a correlation between pLI and the allelic fold change (log2 aFC) associated with the eQTL. We observed that pLI is significantly correlated with aFC (r=-0.07, p<0.01; Fig 3). Ignoring the direction of the change in expression, by using the absolute value of log2 aFC, we identified an even stronger negative correlation between pLI and |aFC| (r=-0.31, p<0.01; Fig 4). Collectively, these results are consistent with the hypothesis that long distance (especially inter-chromosomal) regulatory connections exhibit greater tissue specificity and disease associations^22,33^ because they are enriched for LoF-intolerant gene sets. Thus, the inter-chromosomal regulatory connections potentially highlight novel disease pathways associated with the known cohesinopathies.

**Figure 3.**
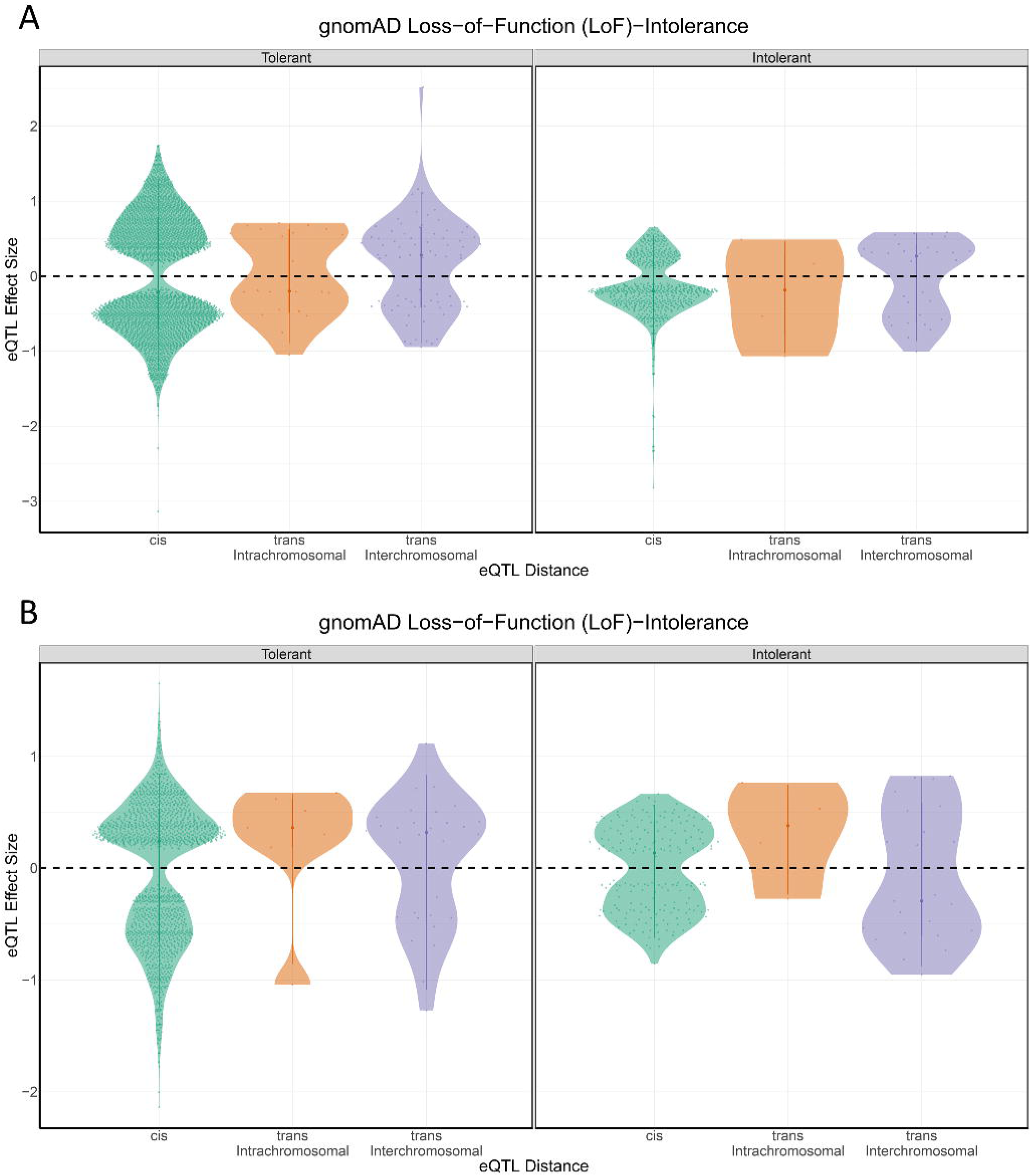
Loss-of-function intolerance shows enrichment for variant-gene eQTL distance. Violin plots of allelic fold change (aFC) across the (a) eQTL-attributed and (b) GWAS-attributed gene sets, pLI and LoF-intolerance (pLI>0.9) are both significantly correlated with aFC.

**Figure 4.**
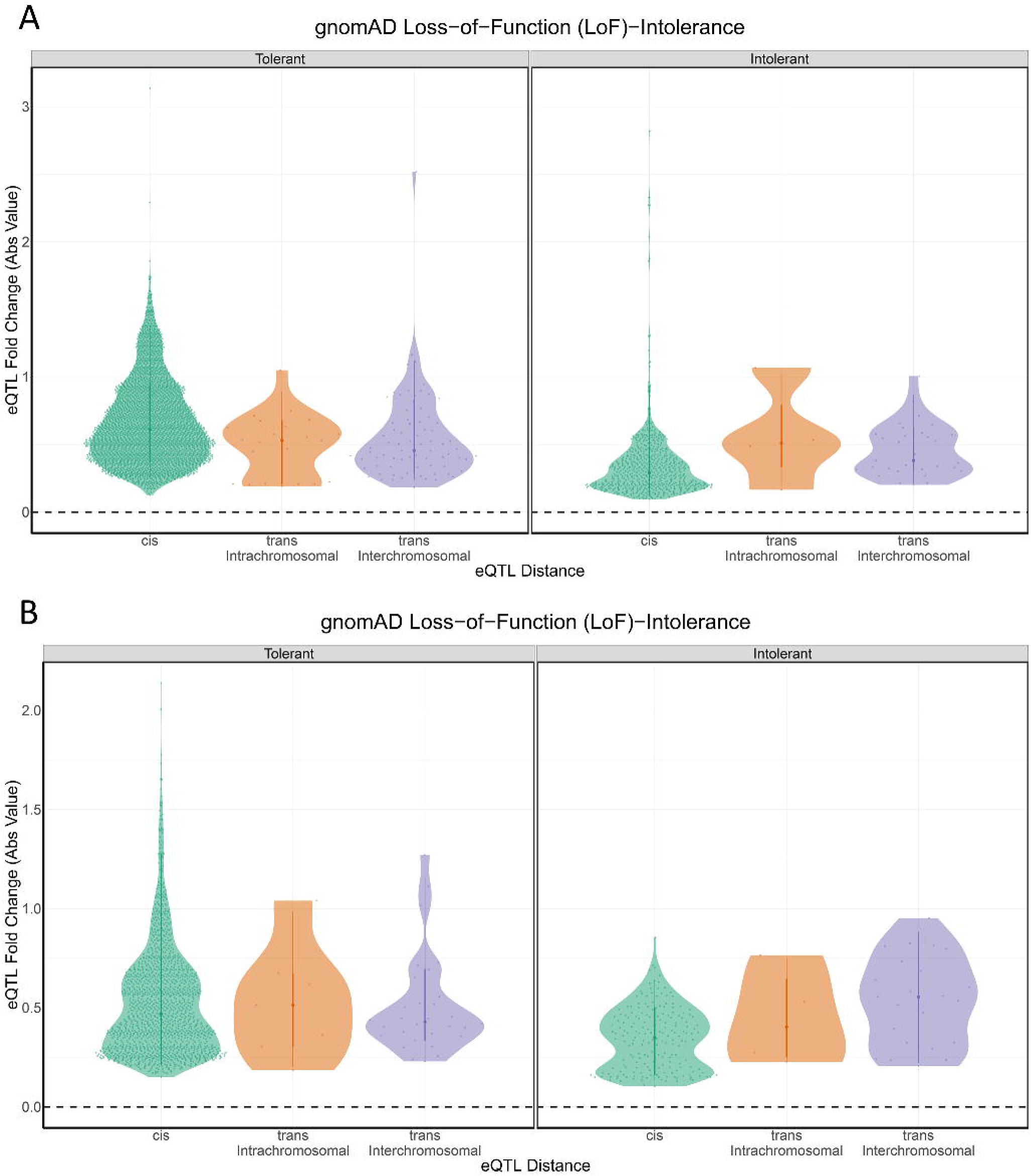
Loss-of-function intolerance shows an effect on eQTL effect size at each variant-gene eQTL distance category. Violin plots of allelic fold change (aFC) across the (a) eQTL-attributed and (b) GWAS-attributed gene sets, pLI and LoF-intolerance (pLI>0.9) are both significantly correlated with aFC as an absolute value.

### Pathway analysis identifies cohesin gene connections with extracellular matrix production and the proteasome

We hypothesized that genes connected to regulatory variants within the cohesin loci might be contributing to disease-related phenotypes. A g:Profiler TRANSFAC analysis identified significant enrichment for target sequences for the EGR1 transcription factor (Table S7). *EGR1* transcription is regulated by stress and growth factor pathways; its binding to DNA is modulated by redox state, and its transcriptional targets include genes involved in extracellular matrix production^34^. g:Profiler enrichment also identified 9 genes as part of the ubiquitin mediated proteolysis pathway (*BIRC3, BIRC6, CUL2, UBE2F, UBE2K, UBE4B, UBR5, CUL3, XIAP*; Tables S7 [red] and S10). The proteasome pathway tags unwanted proteins to be degraded and defects in proteolysis have a causal role in a variety of cancers. Notably, 8 of the 9 (88.9%) genes identified in the Ubiquitin-mediated proteolysis pathway are LoF-intolerant.

### Cohesin gene loci interactions with distant LoF-intolerant genes highlight chromatin-remodelling drug interactions

We hypothesized that the 310 genes we identified would have pharmacokinetic interactions with cancer treatments. Notably, we identified *CYP3A5* as being regulated by a *cis* interaction with a variant 441,888 bp away (intronic, CNPY4). *CYP3A5* encodes a protein within the cytochrome P450 family. Defects in P450 are known to alter cancer treatment outcomes (drug metabolism, KEGG hsa00982). Additionally, we identified a g:Profiler enrichment for the E2F7 transcription factor within our eGenes. E2F7 is up-regulated in response to treatment with doxorubicin or etoposide, topoisomerase 2 blockers^31^. Indeed, many of the 310 genes from the CoDeS3D analysis are targeted by drugs (Table S11). By contrast, for the cohesin genes, the drug-gene interaction database only lists known drug interactions for STAG2 and STAG3^35^. Notably, consistent with our earlier observations, the drug-gene targets we identified include two genes targeted by topoisomerase blockers (*e.g. PAPOLA*, and *XIAP*)^35^, both of which are LoF-intolerant genes.

## Discussion

Through the use of the “Contextualize Developmental SNPs using 3D Information” (CoDeS3D) algorithm^21^, we have leveraged physical proximity (Hi-C) and gene regulatory changes (eQTLs) to reveal how variation in putative enhancers can alter the regulation of cohesin genes and modifier genes. Our analysis has identified 140 eQTLs that link 310 genes associated with the mitotic cohesin ring genes (*SMC1A, SMC3, STAG1, STAG2*, and *RAD21/RAD21*-AS), meiosis-specific cohesin ring genes (*SMC1B, STAG3, REC8*, and *RAD21L1*), cohesin-associated support proteins (*WAPL, NIPBL, PDS5A, PDS5B*, and *MAU2*) and *CTCF*. Collectively, these results form an atlas of functional connections from cohesin genes to proximal and distal genes, some of which reside on different chromosomes to the regulatory elements. These results agrees with previous findings that spatial eQTLs mark hubs of activity across a multi-morbidity atlas^21^.

Only 35% of variant-gene mappings in the GWAS catalogue were supported by spatial *cis*-eQTLs. Therefore, although several GWAS-associated disease SNPs have been linked with cohesin, only a minority have supporting evidence that these variants actually regulate the cohesin genes. As such, our results call into question the validity of many of the previous associations between non-coding genetic variants and the cohesin genes that have been made in the GWAS catalogue.

Previous reports have suggested that cis-eQTL gene sets are depleted of LoF-intolerance genes when compared to similarly sized sets of non-eQTL genes^36^. Similarly, our findings show that the 310 genes regulated by variants in cohesin loci are also enriched for LoF-intolerant genes is notable. The apparent bias in existing studies can be explained by eQTL studies being predominantly focused on nearby (*cis*) variant-gene transcriptional connections and Hi-C studies focusing on local changes in TAD structure, due to technical limitations in the current analysis pipelines.

We revealed that the greater the distance separating the eQTL and target gene, the more likely the target gene was to be LoF-intolerant (*i.e.* over 30% of *trans*-interchromosomal interactions involved LoF-intolerant genes). This finding is consistent with studies that show that long distance connections exhibit greater disease- and tissue-specificity^22,33,37^. The demonstration that the cohesion genes are also LoF-intolerant agrees with their recognised haploinsufficiency in human developmental disease^2,13^. Notably, the genes that were enriched within pathways of pathological importance (*e.g.* 8 of the 9 genes in the ubiquitin-mediated proteolysis pathway gene set) were more likely to be LoF-intolerant. This is consistent with previous findings that eQTL-identified genes are enriched for genome-wide disease heritability, and the subset of eQTL genes with LoF-intolerance are even larger enrichments for genome-wide disease heritability^38^.

The CoDeS3D method identified eQTL links between cohesin genes and other loci not related to cohesin. Two pathways emerged from this analysis. Firstly, spatial eQTL connections with cohesin genes identified an enrichment for genes that are regulated by zinc finger transcription factors including EGR1 and ZNF880. EGR1 positively regulates extracellular matrix (ECM) production. Interestingly, we recently observed widespread dysregulation of extracellular matrix genes upon deletion of cohesin genes in leukaemia cell lines^39^. This supports the idea that regulation of the cohesin complex is tightly associated with ECM production. Additional support for this is derived from the observation that cohesin subunit SMC3 exists in the form of an extracellular chondroitin sulfate proteoglycan known as bamacan^40^. Additionally, in asthma, SMC3 upregulation significantly affected ECM components^41^. The ECM facet of cohesin biology is relatively under-explored and is worthy of further investigation. Secondly, spatial eQTL connections with cohesin genes identified an enrichment for genes that encode effectors of the proteasome pathway. The stability of many cohesin proteins are regulated by the proteasome pathway^1,42,43^, but aside from this, genetic interactions between cohesin genes and proteasome pathway genes remain unexplored.

In conclusion, many studies of mutations focus on the impact of coding-region variation, relying on natural knockouts (especially missense and loss of function variants) to identify gene function. Our analysis highlights what those studies might be missing: sets of co-ordinated genes important to disease but largely intolerant to LoF mutation in healthy individuals. We identified a novel set of genes which are regulated by elements within the cohesin genes. We found that many of the pathways and transcription factor binding sites enriched within these genes were relevant to disease pathways relevant to development and cancer. Moreover, drug-gene interactions further reinforce the importance of these connections to cancer drug treatments and in particular topoisomerase-targeting drugs. As such, our results support recent reports of the importance of long-distance regulation as a key driver of phenotype development^33^.

## Methods

### A large number of GWAS studies have mapped phenotypic variation to cohesin ring gene loci

We searched the GWAS catalogue for SNPs mapped or attributed to mitotic cohesin ring genes (*SMC1A, SMC3, STAG1, STAG2*, and *RAD21*/*RAD21*-*AS*), meiosis-specific cohesin ring genes (*SMC1B, STAG3, REC8*, and *RAD21L1*), cohesin-associated support proteins (*WAPL, NIPBL, PDS5A, PDS5B*, and *MAU2*), and *CTCF* from GWAS studies covering a large assortment of altered phenotypes and pathologies across most tissues in the body (GWAS-attributed).

Genomic positions of SNPs were obtained from dbSNP for human reference hg38.

### Composite Regulatory Impact (*cis*-eQTLs) of Variants in Cohesin Gene Loci

Beyond variants with association to disease, we searched the GTEx catalogue for cis-regulatory variants (variants within 1Mb) that modify the expression of either cohesin ring genes (*SMC1A, SMC1B, SMC3, STAG1, STAG2, STAG3, RAD21, REC8*, and *RAD21L1*), cohesin support genes (*WAPL, NIPBL, PDS5A, PDS5B*, and *MAU2*), or *CTCF* in one or more of 44 tissues across the human body (eQTL-attributed). Unlike GWAS variants, these variants have no inherent association to a phenotype (except the overlaps), as GTEx contains individuals that were relatively healthy prior to mortality. Thus, these variants explain variation in gene expression in a normal, mostly older cohort.

Genomic positions of SNPs were obtained from dbSNP for human reference hg38.

### Identification of SNP-Gene Spatial Relationships

For all GWAS-attributed and eQTL-attributed variants, spatial regulatory connections were identified through genes whose transcript levels depend on the identity of the SNP through both spatial interaction (Hi-C data) plus expression data (expression Quantitative Trail Locus [eQTL]; GTEx v8^45^) using the CoDeS3D algorithm (https://github.com/Genome3d/codes3d-v1)^21,46^. Spatial-eQTL association p-values were adjusted using the Benjamini–Hochberg procedure, and associations with adjusted p-values < 0.05 were deemed spatial eQTL-eGene pairs. Variants not found in the GTEx catalogue or variants with a minor allele frequency below 5% were filtered out due to the sample size of GTEx at each tissue.

To identify SNP locations in the Hi-C data, reference libraries of all possible Hi-C fragment locations were identified through digital digestion of the hg38 human reference genome with the same restriction enzyme employed in preparing the Hi-C libraries (*i.e.* MboI, HindIII). Digestion files contained all possible fragments, from which a SNP library was created, containing all genome fragments containing a SNP. Next, all SNP-containing fragments were queried against the Hi-C databases to find distal fragments of DNA which spatially connect to the SNP-fragment. If the distal fragment contained the coding region of a gene, a SNP-gene spatial connection was confirmed. There was no binning or padding around restriction fragments to obtain gene overlap. To limit technological challenges, gene transcripts for both the spatial and eQTL analyses used the GENCODE transcript model.

Spatial connections were identified from previously generated Hi-C libraries of various origins (Supp Table 12): 1) Cell lines GM12878, HMEC, HUVEC, IMR90, K562, KBM7, HELA, NHEK, and hESC (GEO accession numbers GSE63525, GSE43070, and GSE35156); 2) tissue-specific data from ENCODE sourced from the adrenal gland, bladder, dorsolateral prefrontal cortex, hippocampus, lung, ovary, pancreas, psoas muscle, right ventricle, small bowel, and spleen (GSE87112); and 3) Tissues of neural origin from the Cortical and Germinal plate neurons (GSE77565), Cerebellar astrocytes, Brain vascular pericytes, Brain microvascular endothelial cells, SK-N-MC, and Spinal cord astrocytes (GSE105194, GSE105513, GSE105544, GSE105914, GSE105957), and Neuronal progenitor cells (GSE52457).

### Defining mutationally-constrained genes

The human transcriptome consists of genes with varying levels of redundancy and critical function, resulting in some genes being intolerant to loss-of-function (LoF-intolerant) mutation. This subset of the human transcriptome are posited to also be more intolerant to regulatory perturbation. The gnomAD catalog^32^ lists 19,704 genes and their likelihood of being intolerant to loss-of-function mutations (pLI), resulting in 3,063 LoF-intolerant, 16,134 LoF-tolerant, and 507 undetermined (15.5% LoF-intolerance, defined as pLI ≥ 0.9)^32^. We tested all cohesin, cohesin-associated genes, and those from our analysis (GWAS- and eQTL-attributed) for LoF-intolerance, comparing our *cis* and *trans*-acting eQTL gene lists for enrichment for LoF-intolerance. As pLI is bimodal and non-normally distributed, we tested both pLI raw values as well as pLI grouping (tolerant vs intolerant) for correlation between eQTL effect size (log2 allelic fold change, aFC) and intolerance to disruption (pLI). We considered both aFC and its absolute value (direction of effect ignored), as it has been suggested that the eQTL effect direction is determined by how you define the minor allele within the population, not the actual molecular impact of the eQTL on the cohesin connection. This analysis highlights the significance of long-distance gene regulation on otherwise mutationally-constrained (LoF-intolerant) genes.

### Gene Ontology (GO), Pathway Analysis, and Functional Prediction

All genes from the GWAS-attributed and eQTL-attributed analyses were then annotated for significant biological and functional enrichment using g:Profiler^47^, which includes the Kyoto Encyclopedia of Genes and Genomes (KEGG) Pathway Database (https://www.kegg.jp/kegg/pathway.html) for pathways and TRANSFAC for transcription factor binding enrichment. Finally, we identified drugs that target the genes and related mechanisms through the Drug Gene Interaction database (DGIdb)^35^.

### Annotation algorithms highlight potential for regulatory behaviour

To predict the most phenotypically causal variants within the variant set, we compared variants from the CoDeS3D analysis with several tools which leverage deep learning-based algorithmic frameworks to classify functional relevance from DNA markers including identified chromatin marks (enhancer marks, etc). We used DeepSEA^48^ to predict the chromatin effects of variants to prioritize regulatory variants and PredictSNP2^49^ to summarise estimates of noncoding variants for classification (deleterious or neutral). DeepSEA predicts the chromatin effects of sequence alterations by analysing the epigenetic state of a sequence (transcription factors binding, DNase I sensitivities, and histone marks) across multiple cell types. PredictSNP2 predictions are a consensus score from across five separate prediction tools for variant prioritization: CADD 1.2 and FATHMM-MKL are modelled on SVM prediction, DANN leverages Deep neural networks, FunSeq2 2.1.2 uses a weighted scoring system, and GWAVA 1.0 makes its predictions based on a random forest model.

### URLs

GTEx portal: https://www.gtexportal.org/home/

CoDeS3D pipeline: https://github.com/Genome3d/codes3d-v1

## Acknowledgements

This work was supported by a Royal Society of New Zealand Marsden Grant to JH and JOS (16-UOO-072), and WS was supported by the same grant.

## Competing interests

None declared.

## Figures

**Supplementary Figure 1.**
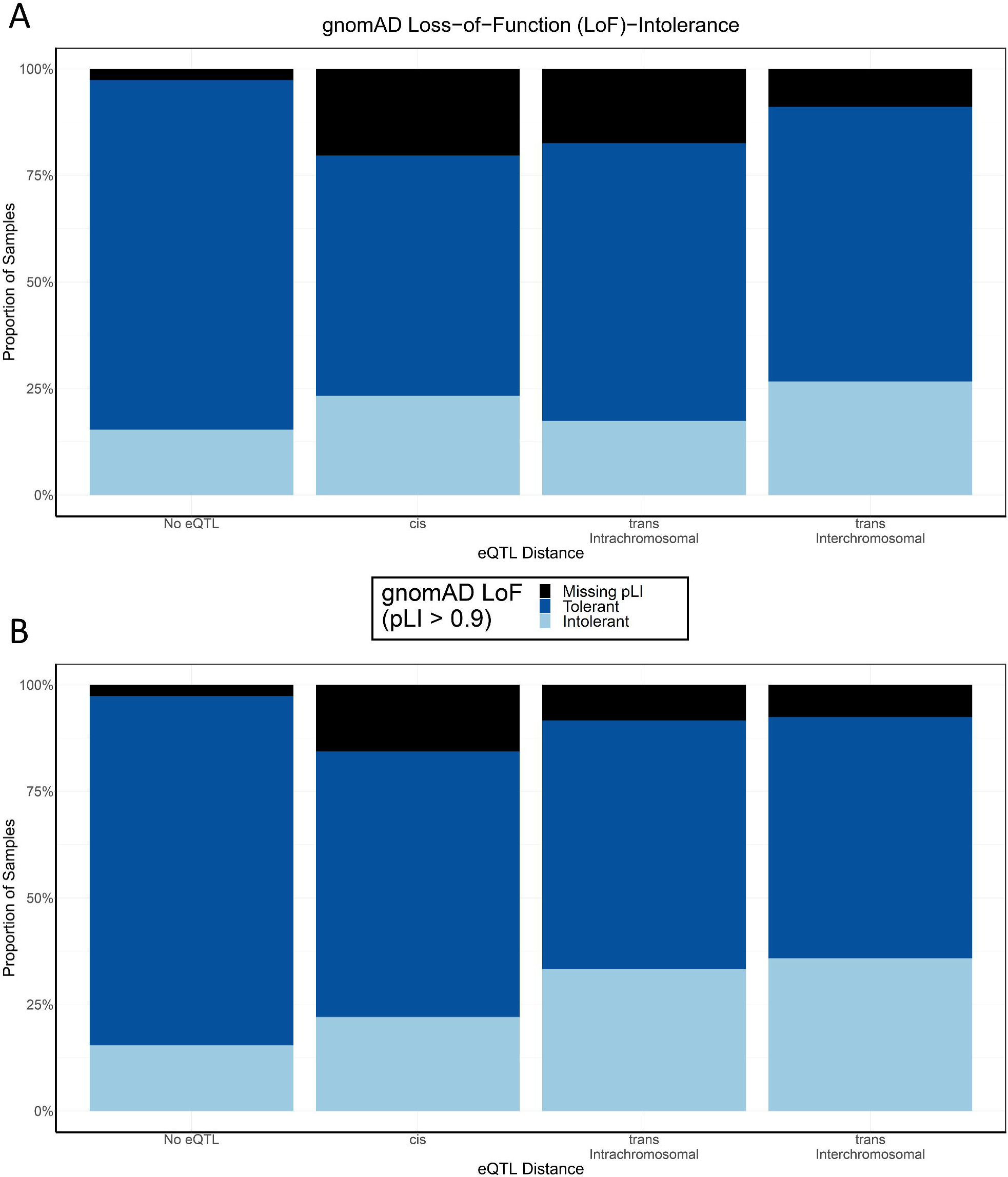
Loss-of-function intolerance (categorical) has the highest prevalence in trans-interchromosomal variant-gene connections. The changing distributions of loss-of-function intolerance across the (a) eQTL-attributed and (b) GWAS-attributed gene highlights genes further away (*trans*) as more LoF-intolerant versus those genes in background (No eQTL) or nearby (*cis*) (21.8% [31 out of 142] *cis*, 23.5% [8 out of 34] *trans*-intrachromosomal, and 30.1% [41 out of 136] *trans*-interchromosomal).

## Tables

Table 1. Number of variants in the eQTL-attributed and GWAS-attributed lists

The eQTL-attributed variant list consists of 187 variants, but after filtering results in 106 variants with significant spatial eQTLs. The GWAS-attributed variant list consists of 106 variants, but after filtering results in 37 variants with significant spatial eQTLs. Overall, the 290 attributed variants results in 140 variants with significant spatial eQTLs.

Table 2. pLI of cohesin genes shows largely mutation intolerance

The main set of genes comprising cohesin and cohesin-support are all loss-of-function intolerant (pLI > 0.9). However, the subset of cohesin genes specific to only meiosis are not loss-of-function intolerant.

Supplementary Table 1. SNP identities for CoDeS3D test for both the eQTL-attributed and GWAS-attributed variant lists

The eQTL-attributed variant list consists of 187 variants, but after filtering results in 106 variants with significant spatial eQTLs. The GWAS-attributed variant list consists of 106 variants, but after filtering results in 37 variants with significant spatial eQTLs. Overall, the 290 attributed variants results in 140 variants with significant spatial eQTLs. Red: variants overlapping each set; Green: variants filtered from CoDeS3D for minor allele frequency (MAF) < 0.05; Blue: variants not in the GTEx variant dictionary (no variant in GTEx).

Supplementary Table 2. CoDeS3D results for GWAS-attributed Cohesin SNPs

CoDeS3D scans the GWAS-attributed variant list (106 variants) for variants with physical proximity to genes that is supported by allele-specific gene expression changes (eQTLs). This analysis found 2188 variant-gene-tissues connections, involving 37 GWAS-attributed variants.

Supplementary Table 3. CoDeS3D results for eQTL-attributed Cohesin SNPs

CoDeS3D scans the eQTL-attributed variant list (187 variants) for variants with physical proximity to genes that is supported by allele-specific gene expression changes (eQTLs). This analysis found 4607 variant-gene-tissues connections, involving 106 eQTL-attributed variants.

Supplementary Table 4. Motifs and Binding Proteins modified by the Variants in this Study (Haploreg v4.1)

We analysed the 209 genetic variants we identified for patterns of histone marks, DNAse accessibility, and protein binding motifs in Haploreg v 4.1. Most of the variants have marks of accessible chromatin (DNAse: 61.3%). In addition, almost all of the variants (93.1%) have at least one of the three: Promoter histone marks (82.8%), Enhancer histone marks (45.1%), proteins binding site (86.8%). Additionally, 16 of the variants modify protein binding or motif predictions in RAD21 (12), SMC3 (4), and/or CTCF (10). Green: protein or motif lists including a cohesin gene.

Supplementary Table 5. DeepSEA estimated effect of the cis-associated and GWAS-associated variants

DeepSEA predicts the chromatin effects of sequence alterations by analysing the epigenetic state of a sequence (transcription factors binding, DNase I sensitivities, and histone marks) across multiple cell types. This results in 28 variants identified as functionally significant.

Supplementary Table 6. PredictSNP2 estimated effect of the cis-associated and GWAS-associated variants

PredictSNP2 predictions are a consensus score from across five separate prediction tools for variant prioritization: CADD 1.2 and FATHMM-MKL are modelled on SVM prediction, DANN leverages Deep neural networks, FunSeq2 2.1.2 uses a weighted scoring system, and GWAVA 1.0 makes its predictions based on a random forest model. This results in 9 variants identified as deleterious.

Supplementary Table 7. Gene list enrichment (g:Profiler 2019)

The 310 genes identified by CoDeS3D were searched for enrichment in various biological processes through g:Profiler, identifying 54 significantly enriched processes. When removing the cohesin genes from the enrichment analysis, 4 processes remain significant (red), including 2 ubiquitin gene ontologies.

Supplementary Table 8. Gene list and g:Profiler enrichment for Meiosis-specific cohesin genes (RAD21L1, REC8, SMC1B, STAG3; g:Profiler 2019)

(a) There are 108 genes functionally connected to variants within the meiosis-specific cohesin loci SMC1B, STAG3, REC8, and RAD21L1. (b) We identified 14 significantly enriched processes, including the gene ontology “male germ cell nucleus” pathway and for TRANSFAC targets for the E2F7 transcription factor, which link to meiosis- and cell-cycle specific mechanisms.

Supplementary Table 9. Loss-of-function intolerance of variant-attriubted genes

The gnomad database identified LoF-intolerant genes (Intolerance defined as pLI ≥ 0.9). Here we show the pLI for the 310 genes identified by spatial eQTLs in our GWAS- and eQTL-attributed variant analysis. Overall, 185 are LoF tolerant, 79 of the 310 genes (25.4%) are LoF-intolerant, and 46 lack pLI.

Supplementary Table 10. Highlights of genes identified within the Ubiquitin mediated KEGG pathway

Within the g:Profiler analysis, two Ubiquitin-mediated KEGG pathways were significant even when removing cohesin from the analysis.. Here we highlight the pathways and their associated genes, eQTL distance, variant-attributed locus, and source of variant-attribution. The KEGG disease pathways identified above are largely identifying LoF-intolerant genes (8 of 9 [88.9%] KEGG Ubiquitin-mediated proteolysis genes).

Supplementary Table 11. Drug-gene interactions with eQTL-attributed and GWAS-attributed eQTL genes (DGIDB v 3.0)

The drug-gene interaction database only lists known drug interactions for STAG2 and STAG3, and only for cancer treatments. However, DGIdb identifies a large number of drug-gene interactions with the 310 genes identified by CoDeS3D. Notably, the drug-gene interaction analysis identifies two topoisomerase targets: Camptothecin (STAG2 interaction, a Topoisomerase-I inhibitor used in cancer treatments) and etoposide (PAPOLA and XIAP interactions, which inhibits the topoisomerase II enzyme).

Supplementary Table 12. Hi-C datasets used in this study

Spatial connections were identified from previously generated Hi-C libraries of various origins: 1) Cell lines GM12878, HMEC, HUVEC, IMR90, K562, KBM7, HELA, NHEK, and hESC (GEO accession numbers GSE63525, GSE43070, and GSE35156); 2) tissue-specific data from ENCODE sourced from the adrenal gland, bladder, dorsolateral prefrontal cortex, hippocampus, lung, ovary, pancreas, psoas muscle, right ventricle, small bowel, and spleen (GSE87112); and 3) Tissues of neural origin from the Cortical and Germinal plate neurons (GSE77565), Cerebellar astrocytes, Brain vascular pericytes, Brain microvascular endothelial cells, SK-N-MC, and Spinal cord astrocytes (GSE105194, GSE105513, GSE105544, GSE105914, GSE105957), and Neuronal progenitor cells (GSE52457).

## Notes

### Competing Interest Statement

The authors have declared no competing interest.

